# From Micro to Macro: Avian Chromosome Evolution is Dominated by Natural Selection

**DOI:** 10.1101/2024.11.29.626112

**Authors:** James M. Alfieri, Kevin Bolwerk, Zhaobo Hu, Heath Blackmon

## Abstract

Birds display striking variation in chromosome number, defying the traditional view of highly conserved avian karyotypes. However, the evolutionary drivers of this variability remain unclear. To address this, we fit probabilistic models of chromosome number evolution across birds, enabling us to estimate rates of evolution for total chromosome number and the number of microchromosomes and macrochromosomes while simultaneously accounting for the impact of other evolving traits. Our analyses revealed higher rates of chromosome fusion than fission across all bird lineages. Notably, much of this signal was driven by Passeriformes, where migratory species showed a particularly strong bias towards fusions compared to sedentary counterparts. Furthermore, a robust correlation between the rearrangement rates of microchromosomes and macrochromosomes suggests that genome-wide processes drive rates of structural evolution. Additionally, we found that lineages with larger population sizes exhibited higher rates of both fusion and fission, indicating that positive selection plays a dominant role in driving divergence in chromosome number. Our findings illuminate the evolutionary dynamics of avian karyotypes and highlight that, while the fitness effects of random structural mutations are often deleterious, beneficial mutations may dominate karyotype divergence in some clades.

## Introduction

Historically, avian karyotypes have been characterized as highly conserved with few exceptions, as most birds have a diploid number around 80, consisting of both microchromosomes and macrochromosomes (Griffin et al. 2007; Ellegren 2010). Indeed, bird genomes have been shown to have strikingly fewer intrachromosomal rearrangements than is typical in other clades (e.g., mammals and insects) (Li et al. 2021; Alfieri et al. 2023). However, recent work challenges the view that avian karyotypes are largely static. Degrandi et al. (2020) reported a wide range of diploid counts, from as few as 40 to over 140 chromosomes (**Figure 1**), highlighting substantial variation in chromosome number. This diversity is potentially significant, as chromosome architecture can influence recombination frequency, genetic linkage, hybrid compatibility, sex-determining mechanisms, and evolutionary rates (Stebbins 1958; White 1973; Britton-Davidian et al. 1990; Blackmon and Demuth 2015; Blackmon et al. 2015). Given the profound impacts that karyotype evolution can have and the vast dataset available (i.e., over 1,087 reported avian karyotypes), it is surprising that we have yet to determine whether chromosome number evolution in birds is primarily driven by natural selection or genetic drift and by extension whether extant variation in chromosome number has been generated through the fixation of deleterious, neutral, or beneficial mutations.

**Figure 1.**
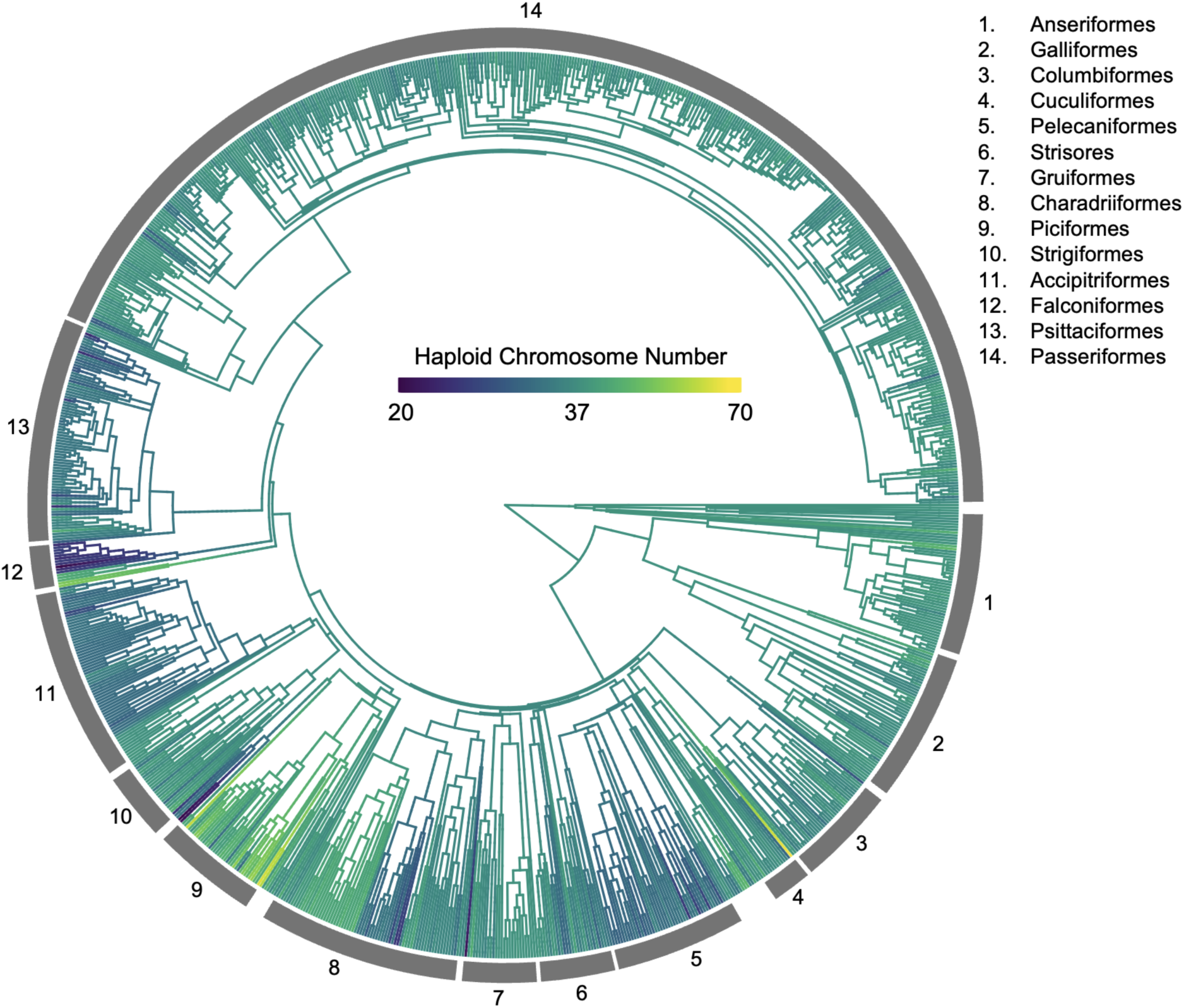
Haploid Chromosome Numbers Across Birds. Each branch on the phylogeny is painted to indicate chromosome number and the color ramp is on a log scale to ease visualization. The bands on the outside of the phylogeny indicate major orders.

Theoretical investigations have played a crucial role in shaping our understanding of chromosome number evolution. Early models, such as those proposed by Lande (1979), explored the conditions under which chromosomal rearrangements could become fixed in populations. These models emphasized the role of genetic drift, particularly in small populations, where the fixation of deleterious chromosomal mutations might occur despite their initial fitness costs. However, these theoretical approaches also have significant weaknesses. The assumption that chromosomal mutations are largely underdominant (i.e., deleterious when heterozygous) limits the applicability of these models, particularly in cases where chromosomal changes may have neutral or even beneficial effects.

Empirical studies have provided valuable, albeit limited, insights into the fitness effects of structural mutations. Radiation induced chromosomal mutations in both *Drosophila melanogaster* and *Tribolium castaneum* have allowed us to measure the fitness effect of a range of structural mutations but it is unclear whether the fitness effects of induced mutations is similar to the fitness effects of naturally occurring mutations (Muller 1930; Sokoloff 1977). Similarly, controlled crosses between karyotypically distinct strains or species have provided additional measures of the fitness of individuals heterozygous for chromosomal mutations (Ratomponirina 1988; Britton-Davidian et al. 1990). However, these crosses typically also involve genic divergence between the strains or species being crossed making it at best difficult to isolate the effects of structural variation alone. Despite these limitations, a consensus emerged that most chromosomal mutations are likely deleterious, particularly in heterozygous individuals (King 1995).

With this base assumption that most chromosomal mutations are deleterious, researchers began attempting to understand the characteristics that could lead to accelerated rates of chromosome number evolution in lineages. Although formal phylogenetic comparative methods were unavailable at the time, studies comparing evolutionary rates across placental mammals, other vertebrates, and mollusks (Wilson et al. 1975), among mammalian clades (Bush et al. 1977), and among beetle families (Petitpierre 1987) all suggested that clades with smaller effective population sizes had higher rates of evolution. Recent advances in comparative phylogenetic methods have provided new opportunities to investigate the forces shaping chromosome number evolution and for statistical tests of a range of hypotheses (Mayrose et al. 2010; Glick and Mayrose 2014; Zenil-Ferguson et al. 2017; Freyman and Höhna 2018; Blackmon et al. 2019). These tools allow researchers to explicitly test hypotheses while controlling for phylogenetic history. Empirical studies leveraging these comparative approaches have begun to reveal patterns across large clades that were not accessible in the past.

Some clades (i.e., Carnivores and Coleoptera) show strong support for a pattern where lineages with reduced effective population size have increased rates of chromosome number evolution (Blackmon et al. 2024; Jonika et al. 2024). These results suggest that karyotype diversity in these lineages may be generated primarily through the fixation of deleterious mutations that are fixed by genetic drift. Outside of Carnivores most mammalian lineages show a pattern that is consistent with female meiotic drive as the primary driving force in generating karyotypic diversity (Blackmon et al. 2019). While probabilistic models of chromosome number evolution have not (to our knowledge) been applied to birds as a whole, some studies have looked at rates of structural evolution in subclades. For instance, the rate of inversions in island and mainland Estrildid finches was compared and demonstrated that inversions fix more frequently in mainland species suggesting that these inversion are fixed by natural selection (Hooper and Price 2015).

Bird karyotypes are notable for their combination of macrochromosomes and microchromosomes. This bimodal structure has been largely conserved since the emergence of modern birds, with cytogenetic and comparative genomic evidence suggesting that the ancestral avian karyotype resembled that of present-day species, such as chickens (Griffin et al. 2007; O’Connor et al. 2018; Zhang 2018). Microchromosomes differ from macrochromosomes in several key ways: they are much smaller (Pichugin et al. 2001), typically more gene-dense (Smith et al. 2000), and exhibit higher rates of recombination (Rodionov et al. 1992). These characteristics lead to an expectation that microchromosomes should have higher evolutionary rates of rearrangements than macrochromosomes. However, microchromosomes show remarkable conservation in gene content and organization across extremely diverged avian species (Damas et al. 2018; Waters et al. 2021), suggesting that rates of microchromosomes and macrochromosomes are decoupled. This conservation suggests purifying selection against structural rearrangements in microchromosomes. In contrast, macrochromosomes exhibit greater variability, suggesting a higher rate of structural rearrangements, particularly in some specific lineages (O’Connor et al. 2019, 2024; Kretschmer et al. 2021).

In this study, we aim to clarify whether chromosome number evolution in birds is primarily driven by genetic drift or natural selection and determine if the tempo and mode of evolution in microchromosomes and macrochromosomes is coupled or decoupled. By leveraging extensive comparative analyses of avian karyotypes, we demonstrate that species with larger population sizes have elevated rates of chromosome fission and fusions, suggesting natural selection as a key force in shaping karyotype dynamics. Additionally, our results indicate that microchromosomes and macrochromosomes have coupled evolutionary rates of fissions and fusions. These findings provide insights into the balance of adaptive and stochastic processes driving genomic features in birds.

## Methods

### Population Size

We had two approaches for investigating the role of population size on chromosome number evolution. The first approach was to characterize orders (with greater than 20 species) into population size groups (small, medium, and large) based on three traits hypothesized to be associated with effective population size (N_e_). The traits used were body mass; small body mass associated with lower N_e_ (Eo et al. 2011), range size; small range size associated with lower N_e_ (Gaston and Blackburn 1996), and trophic level; with expected N_e_ lowest in carnivores, intermediate in omnivores and highest in herbivores (Brüniche-Olsen et al. 2021). We obtained this trait data from Tobias et al. (2022). For each of these traits we separated species into the lowest quartile, the middle two quartiles, or the upper quartile. Each order was then assigned a value of −1, 0, or 1 respectively based on which of these quartiles most species fell into.

For body mass, the lowest quartile of masses was less than 21.40 grams. The second group contained the middle two quartiles, and the third group contained the highest quartile of masses that were greater than 460 grams. For range size, the lowest quartile of range sizes was less than 1.5 million sq km. The second group contained the middle two quartiles, and the third group contained the highest quartile of range sizes greater than 10.35 million sq km. We classified the trophic levels of species into 3 groups (i.e., herbivores, omnivores, and carnivores). Similar to the previous two traits each order was assigned a value of −1, 0, or 1 depending on which trophic level was most common in the order. These values were then summed for each order leading to population size indices that ranged from −1 to 2 (**Table 1**). We treated orders as small N_e_ if their indices were 1 or 2. Orders with indices of 0 were considered medium N_e_. Finally, orders with indices of −1 were treated as large N_e_.

**Table 1.**
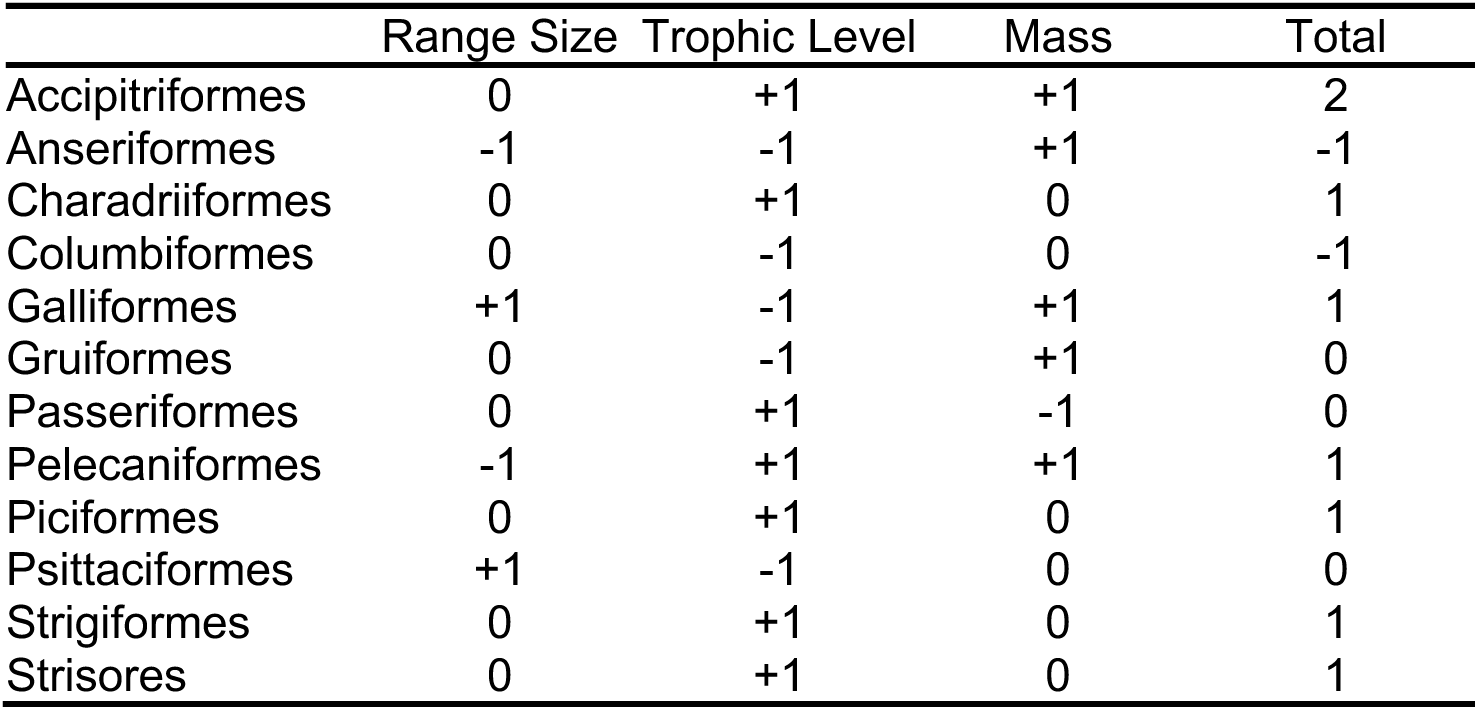
Effective population size scoring scheme. Positive values represent small population size species, while negative numbers represent large population size species.

|  | Range Size | Trophic Level | Mass | Total |
| --- | --- | --- | --- | --- |
| Accipitriformes | 0 | +1 | +1 | 2 |
| Anseriformes | -1 | -1 | +1 | -1 |
| Charadriiformes | 0 | +1 | 0 | 1 |
| Columbiformes | 0 | -1 | 0 | -1 |
| Galliformes | +1 | -1 | +1 | 1 |
| Gruiformes | 0 | -1 | +1 | 0 |
| Passeriformes | 0 | +1 | -1 | 0 |
| Pelecaniformes | -1 | +1 | +1 | 1 |
| Piciformes | 0 | +1 | 0 | 1 |
| Psittaciformes | +1 | -1 | 0 | 0 |
| Strigiformes | 0 | +1 | 0 | 1 |
| Strisores | 0 | +1 | 0 | 1 |

We used the R package ChromePlus to construct Markov models of chromosome evolution allowing for two mechanisms of chromosome number change: fusions and fissions (Blackmon et al. 2019, 2023; R Core Team 2024). We fit a model for each order with more than 20 species. We used chromosome number data from the Bird Chromosome Database (Degrandi et al. 2020). We used a recent phylogeny containing most bird species (McTavish et al. 2024). We fit the models in a Bayesian framework using R package diversitree (FitzJohn 2012). Each Markov Chain Monte Carlo (MCMC) run was initialized with parameter values drawn from a uniform distribution from 0 to 1 and uniform priors. MCMCs were run for 4000 generations. Although all MCMCs appeared to reach convergence at 1000 generations, we conservatively discarded the first 2000 generations as burn-in. All reported rates are lambda parameters for exponential distributions describing the expected waiting time for a transition to occur and are in units of millions of years. From these estimated rates, we tested for phylogenetic signal and performed a Spearman rank correlation analysis between the population size score and the mean rate of fissions and fusions (Revell 2012).

Our second approach for investigating the impact of population size on rates of chromosome number evolution was to utilize global abundance estimates (species census population size) for every species in our dataset, regardless of order representation. We obtained global abundance estimates from Callaghan et al. (2021). Species in the lowest quartile of abundance estimates (< 1395806) were considered small population size species, while species in the highest quartile of abundance estimates (> 16082707) were considered large population size species. After subsetting for species in these quartiles, we incorporated the presence of this binary trait (population size) and transitions between states into our model of chromosome evolution (Blackmon et al. 2023). Each MCMC run was initialized with parameter values drawn from a uniform distribution from 0 to 1 and uniform priors. The MCMC was run for 10000 generations. Although the MCMC appeared to reach convergence at 1000 generations, we conservatively discarded the first 2000 generations as burn-in.

### Microchromosome vs. Macrochromosome Rates of Evolution

We searched the literature compiled in the Bird Chromosome Database for instances where microchromosomes were differentiated from macrochromosomes (Degrandi et al. 2020). This involved searching 353 references containing 1,562 karyotype descriptions of 1086 species. We only included data in which the authors had explicitly provided the number of microchromosomes and macrochromosomes in the karyotype. Authors were not consistent in their definitions of microchromosomes or the heuristics they chose to use. Some authors defined microchromosomes with relative length cut-offs (Hassan 1998; Ebied et al. 2005), while others defined them using cytological and staining techniques (Kaul and Ansari 1978; de Oliveira Furo et al. 2017), and still others differentiate them based on a notable size difference (Goldschmidt et al. 1997). Despite these variations in definitions, the categorization into microchromosomes and macrochromosomes has been strongly supported by the genomics revolution. Specifically, microchromosomes have been shown to have a size distribution independent of macrochromosomes (Burt 2002; International Chicken Genome Sequencing Consortium 2004), elevated GC and gene content (International Chicken Genome Sequencing Consortium 2004; Ellegren 2010), elevated recombination rates (even after controlling for their small size) (Griffin et al. 2007; Skinner and Griffin 2012), and evolutionary conservation across broad lineages (Ellegren 2010; Romanov et al. 2014).

We estimated the rates of fissions and fusions for microchromosomes and macrochromosomes separately. We ran models of chromosome evolution separately for orders with more than 10 species with both microchromosome and macrochromosome counts (N = 8 orders). Each MCMC run was initialized with parameter values drawn from a uniform distribution from 0 to 1. Initial runs showed that the MCMC occasionally failed to mix well and instead explored regions of parameter space that were characterized by unrealistically high rates. To remedy this we used a weak broad exponential prior with a rate parameter of 0.1. We then ran each MCMC for 4000 generations, and discarded the first 2000 as burn-in. We used R package phytools to test for phylogenetic signal (Revell 2012). We then performed a Spearman Rank Correlation test to determine if the mean rate of microchromosome evolution was correlated with the mean rate of macrochromosome evolution.

## Results

We first estimated fission and fusion rates on all birds which had chromosome number and phylogenetic information (N=1027). We found that fusion rates were slightly higher than fission rates. The mean of the fission rate estimates was 0.12 (95% credible interval 0.09 to 0.15). The mean of the fusion rate estimates was 0.20 (95% credible interval 0.16 to 0.23).

### Population Size

Our first approach to investigate the impact of population size on rates of evolution was to characterize orders (with greater than 20 species) into population size groups (small, medium, and large) based on traits hypothesized to affect effective population size (body mass, range size, and trophic level). We had 12 orders with species counts greater than 20: Accipitriformes, Anseriformes, Charadriiformes, Columbiformes, Galliformes, Gruiformes, Passeriformes, Pelecaniformes, Piciformes, Psittaciformes, Strigiformes, and Strisores. In total, 892 species were included in these order-level models (**Table 2**; **Figure 2; Supplemental 1**). Our original models for Piciformes yielded extremely variable rates (mean rate = 2.43/MY, 95% Credible Interval: 0.71 - 6.21). It has been well documented that many comparative methods can be heavily influenced by extreme signals in a very small portion of the phylogeny (Rabosky and Goldberg 2015). To determine if this was the case, we used the tipRate function in the R package evobiR (Jonika et al. 2023) and found one species, *Pteroglossus aracari*, that exhibited recent, drastic reduction in chromosome number (Haploid chromosome number = 31) compared to its closest relative diverged 5.76 MYA, *Pterroglossus castanotis* (Haploid chromosome number = 43). To determine if this tip was leading to our elevated rate estimate in Piciformes, we removed this outlier and repeated our analysis. Even after removal of this taxa Piciformes still had the highest rates of fission and fusion across all examined orders. However, we found that removal of this species led to a 3-fold reduction in the mean rate estimate for this order. In an effort to focus on broadly supported rates of evolution we pruned this taxa from further analyses.

**Figure 2.**
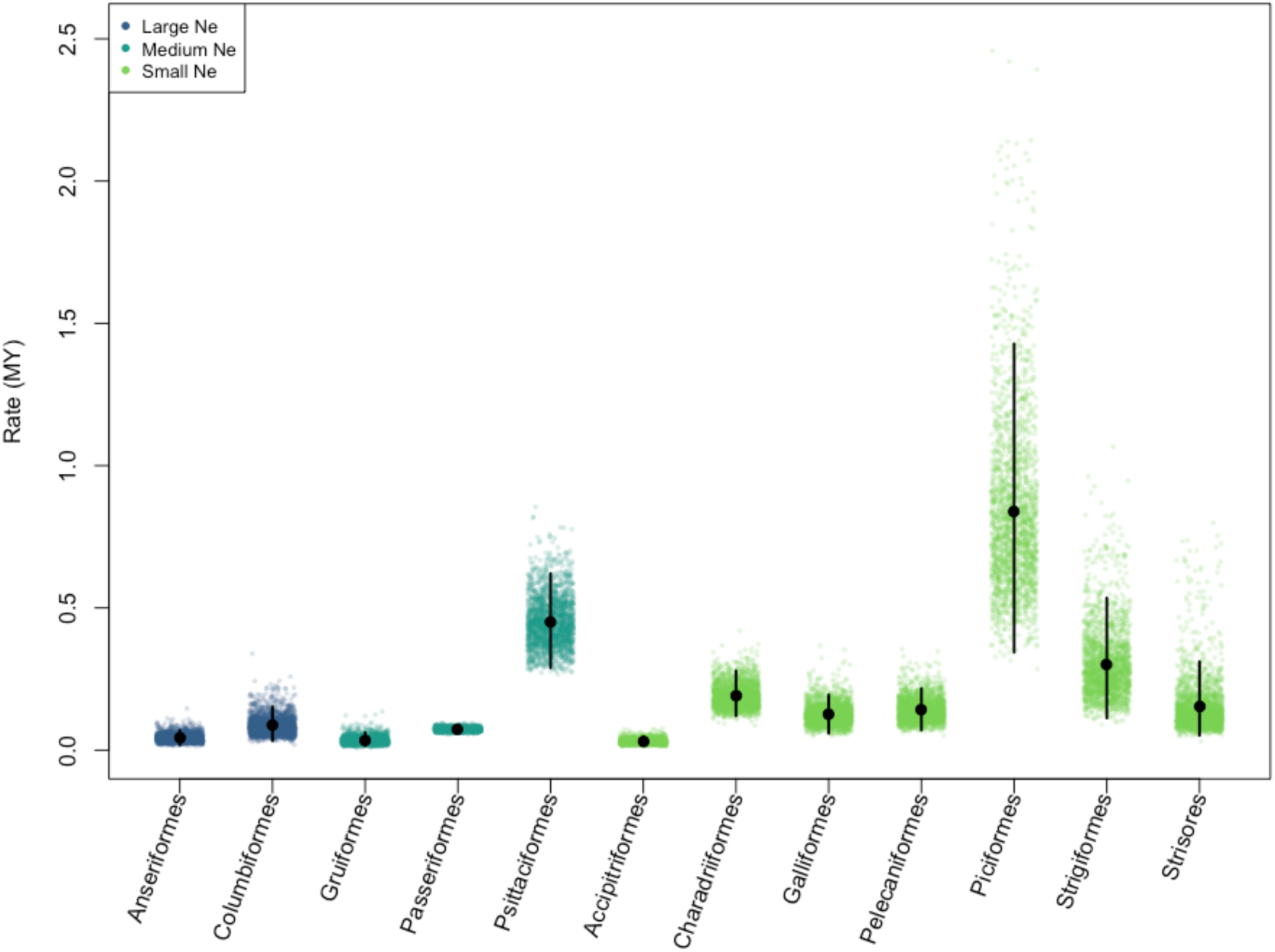
Mean rates of chromosome evolution across bird orders. Each clade displays 2000 posterior samples of mean rate estimates. The posterior mean is marked with a black circle, and the 95% credible interval is shown with a vertical black line. Colors represent the N_e_ class for each clade. The orders are in order of decreasing population size score.

**Table 2.**
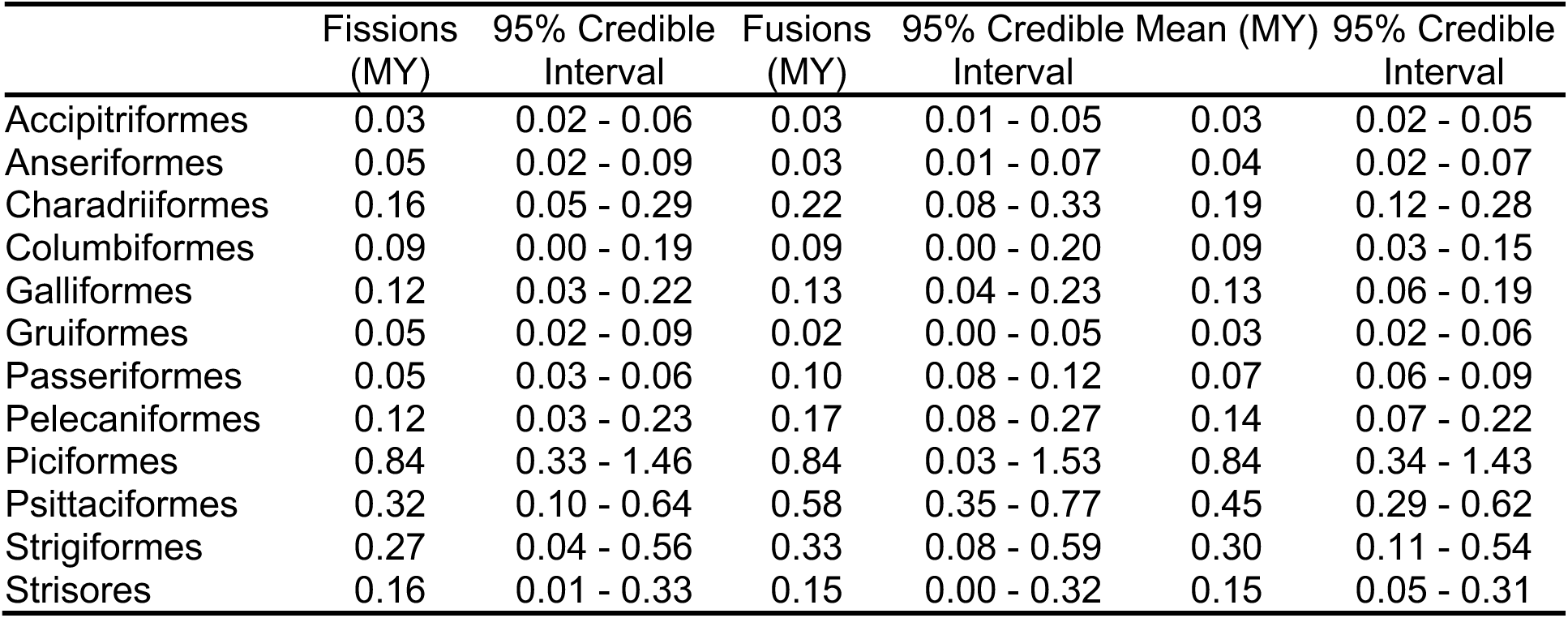
Order level rates of chromosome evolution. For each of 12 orders columns indicate the fission, fusion, and mean rates as well as credible intervals for each.

We did not find any phylogenetic signal in the residuals of the relationship between mean rate and the N_e_ classification (K = 0.87, p = 0.767). There was no linear relationship between N_e_ classification and mean rate (p = 0.41), and there was no correlation (Spearman’s Rho = 0.39, S = 173.26, p = 0.205).

Our second approach was to utilize global abundance estimates (species census population size) for every species in our dataset regardless of order. We had 480 birds in our analysis (240 in the small population size state and 240 in the large population size state). Several small orders were monomorphic for population size, 10 orders were completely classified as small pop size (N = 29) while 6 orders were completely classified as large pop size (N = 8).

The mean fission and fusion rates for large population size lineages was 0.36 and 0.35 respectively (95% Credible Interval for both parameters was: 0.28 - 0.43). The mean fission and fusion rates for small population size lineages was 0.02 and 0.01 respectively (95% Credible Intervals of 0.01 - 0.03 and 0.002 - 0.03 respectively). We computed a mean rate difference statistic, denoted as ΔR_*X*_, where ***X*** is the model parameter of interest (fissions or fusions). For each post-burnin sample, we calculated ΔR_*X*_ as follows: ΔR_*X*_ = ***X***_small population_ - ***X***_large population_. We then assessed the 95% credible interval for ΔR_*X*_. If this interval is entirely below zero it provides strong evidence that the fission and/or fusion rates are higher in clades with large population sizes. Conversely, if the credible interval includes zero, it suggests little to no evidence for a difference in rates between clades with small and large population sizes. We estimated the mean ΔR for fissions to be −0.34 (95% Credible Interval: −0.41 to −0.26), and the mean ΔR for fusions to be −0.34 (95% Credible Interval: −0.42 to −0.26). As the credible intervals do not overlap zero, we infer that large population size species have higher rates of chromosomal fissions and fusions (**Figure 3**).

**Figure 3.**
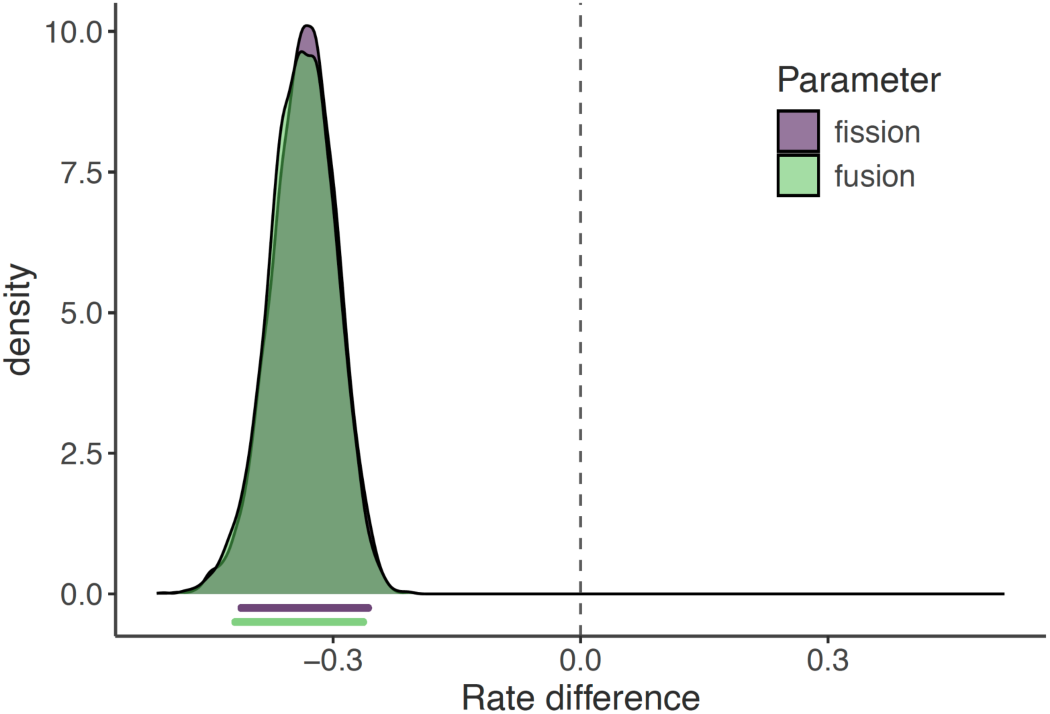
Chromosome evolution in small population size and large population size lineages. Each curve shows the distribution of the rate difference in small population size and large population size lineages. The bars below the curve indicate the 95% credible interval for the rate differences.

To assess whether our findings were disproportionately influenced by a few species with extreme chromosome counts, we conducted a secondary analysis. Following Jonika et al. (2024), we used the GetTipRates function in R package evobiR (Jonika et al. 2023) to calculate the difference between an extant species’ chromosome number and the most probable chromosome number of the immediate ancestor. Species with exceptionally high tip rates could reduce the generalizability of our results to broader clades. To address this, we repeated our analysis–rate estimation with MCMCs and ΔR_*X*_ calculation–excluding species with tip rates greater than 0. This analysis revealed a similar pattern, with large population species exhibiting faster rates than small population species, although the magnitude of the difference was somewhat reduced (**Supplemental 2**).

### Post-hoc Analysis on Passeriformes Migratory Behavior

We were surprised by how Passeriformes was the only bird order that had elevated rates of fusions relative to fissions. We hypothesized that this rate difference might ultimately be due to differences in migratory behavior among Passeriformes. Specifically, we reasoned that migratory birds may more frequently be under selection for coadapted gene complexes important in adaptation to diverse environments (Stebbins 1971). To investigate this we performed a post-hoc analysis. We obtained migration data from Tobias et al. (2022). We incorporated the presence of this binary trait (migratory or sedentary) and transitions between states into our model of chromosome evolution (Blackmon et al. 2023). Each MCMC run was initialized with parameter values drawn from a uniform distribution from 0 to 1 and uniform priors. The MCMC was run for 10000 generations. Although the MCMC appeared to reach convergence at 1000 generations, we conservatively discarded the first 2000 generations as burn-in. We estimated ΔR statistics for fusion and fission rates by subtracting the rate inferred in migratory lineages from the rate inferred in sedentary lineages. The mean of the ΔR_fusion_ statistic was −0.08 with a credible interval of −0.14 to −0.03. The mean of the ΔR_fission_ statistic was −0.02 with a credible interval of −0.07 to 0.02. These results suggest that migratory Passeriformes have higher rates of fusions than sedentary Passeriformes, but equal rates of fissions (**Supplemental 3**).

### Microchromosome vs. Macrochromosome Rates

We had 345 species which had both phylogenetic and microchromosome information. These 345 species represented 24 orders. Across these orders, we found a significant negative relationship between the number of microchromosomes and number of macrochromosomes (**Figure 4**).

**Figure 4.**
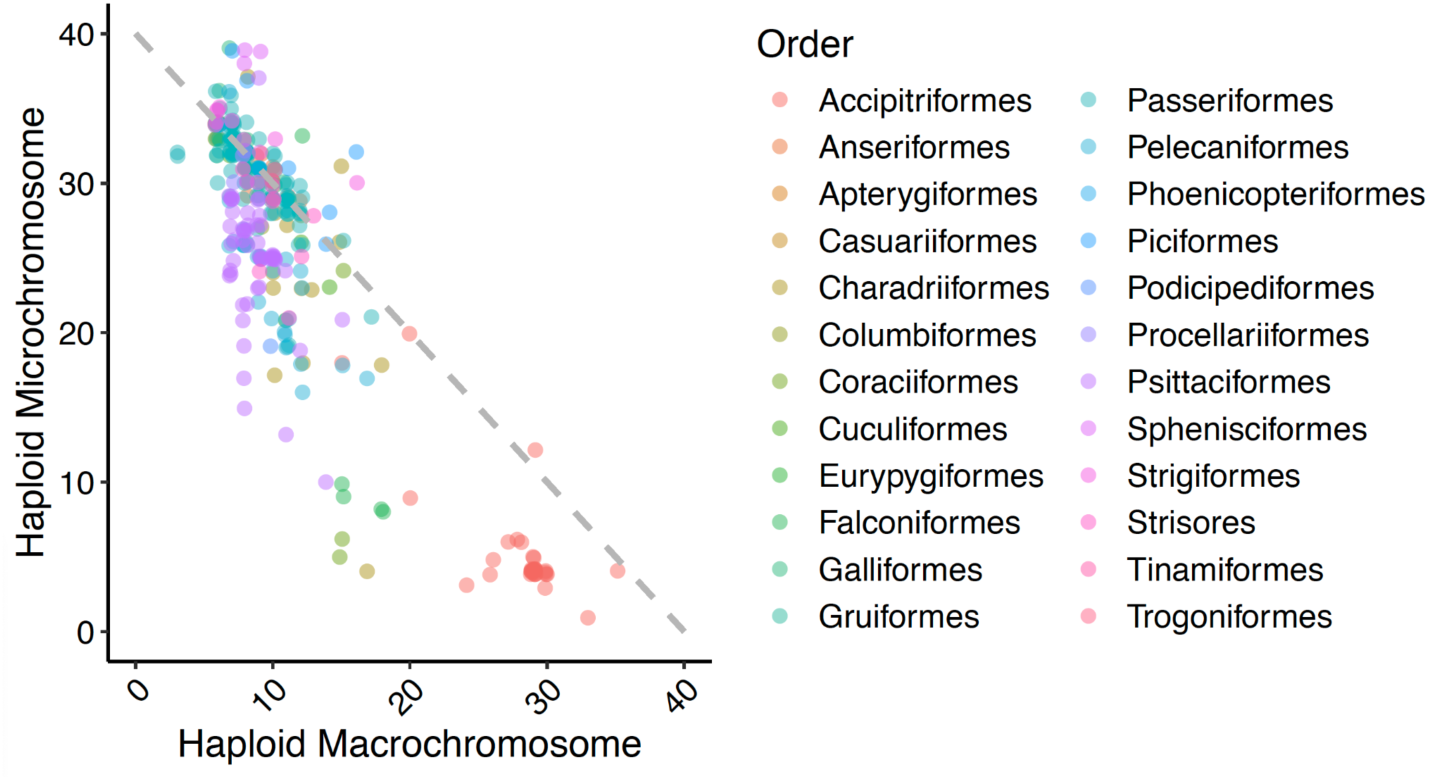
Haploid count of microchromosomes and macrochromosomes. The data plotted are all 339 species for which both microchromosome and macrochromosome counts have been reported. The points are slightly jittered to minimize complete overlaps, and colors represent the order in which the species belongs. The grey dashed diagonal indicates a sum of 40 which is the modal value for our dataset.

We were interested in determining whether rates of microchromosomes and macrochromosomes evolution were coupled. For our analysis, we retained orders with more than 10 species (N = 8): Accipitriformes, Anseriformes, Charadriiformes, Galliformes, Gruiformes, Passeriformes, Pelecaniformes, and Psittaciformes. For each order we fit a simple model of chromosome evolution where the chromosome number was the haploid count of macrochromosomes or microchromosomes. Each MCMC was initiated with parameter values drawn from a uniform distribution and run for 4000 generations with 2000 generations discarded as burnin. We did not find a significant phylogenetic signal of the relationship between microchromosome and macrochromosome evolutionary rates across these orders (K = 0.859, p = 0.643). To determine whether rates for microchromosomes and macrochromosomes were coupled we calculated Spearman’s correlation coefficient for the mean rate of microchromosome and macrochromosome evolution across the eight included orders. This process was repeated for each of our 2000 samples from the posterior distribution of rate estimates. The credible interval of the correlation coefficient ranged from 0.02 to 0.88. This suggests that rates of microchromosome and macrochromosome evolution are coupled (**Figure 5**).

**Figure 5.**
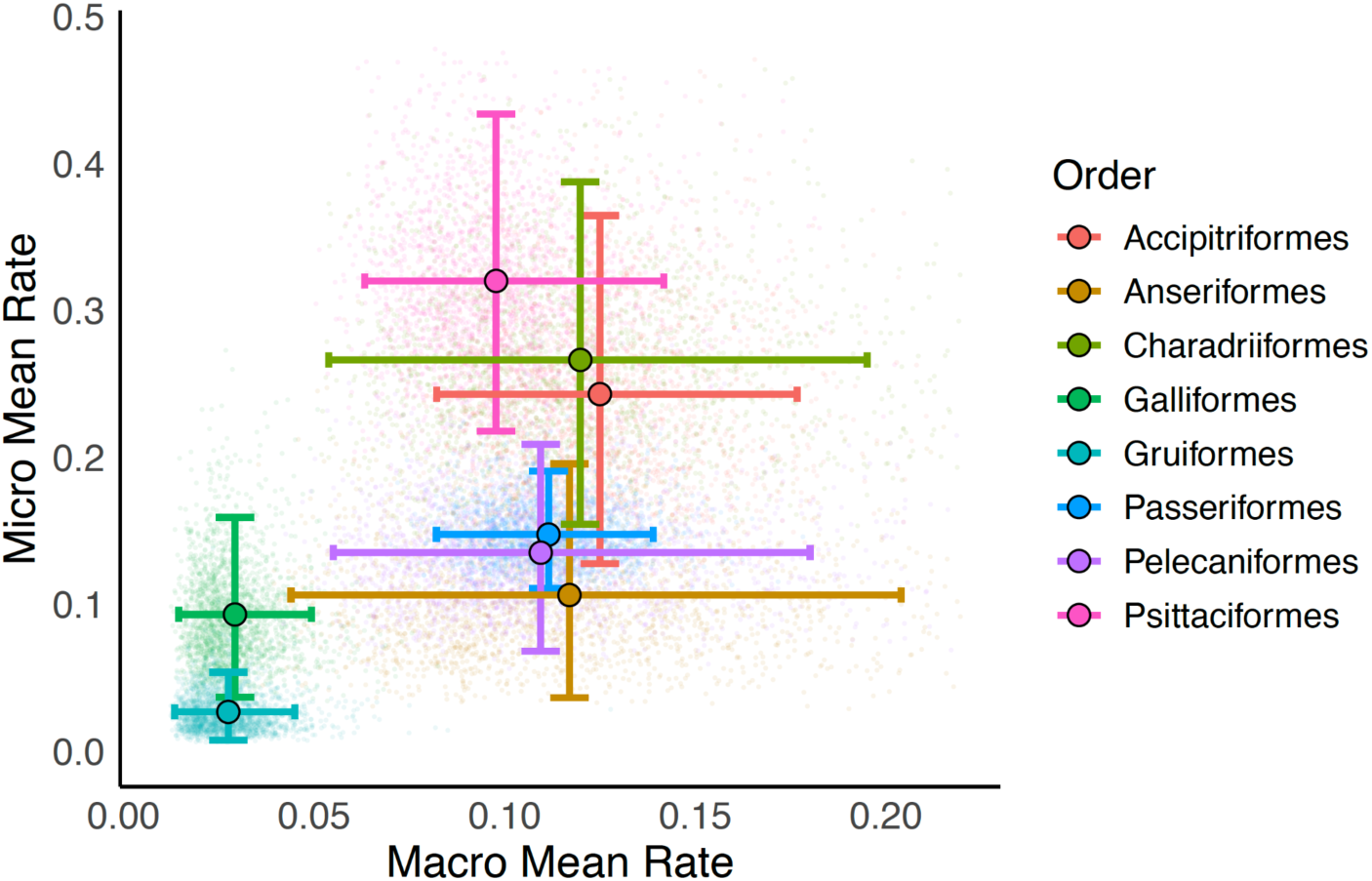
Rates of evolution in microchromosomes and macrochromosomes. The horizontal dimension indicates the mean of the fusion and fission rate in macrochromosomes while the vertical dimension indicates the mean of the fission and fusion rate in microchromosomes. Error bars indicate the 95% credible interval. All elements are colored to reflect the order. The plot is cropped for visualization purposes and fails to include 165 (approximately 1%) of the sampled macrochromosome rates that are well beyond the credible interval.

## Discussion

### Selection vs Drift

A central goal of our study was to determine whether natural selection or genetic drift dominates the evolution of chromosome number in birds. By extension this also allows us to determine the average fitness effect of mutations that have led to divergence in chromosome number among birds (Mackintosh et al. 2023). Briefly, if lineages that have large N_e_ have higher rates of evolution than those with small N_e_, we can infer that selection dominates chromosome number evolution and that these mutations must be on average beneficial (Qumsiyeh and Handal 2022). In contrast, if lineages that have small N_e_ have higher rates of evolution than those with large N_e_ we can infer that drift must dominate chromosome number evolution and that these mutations must be on average deleterious (Wilson et al. 1975; Bush et al. 1977). Finally, if we find no significant differences in the rates of evolution among small and large N_e_ lineages then we would infer that on average these mutations are effectively neutral.

We did not find a significant relationship between population size–measured using the proxies body mass, range size, and trophic level–and chromosomal fission and fusion rates when categorizing entire orders into homogenous population size groups. This lack of a relationship could stem from several factors. First, our proxies may not accurately reflect effective population size. Additionally, characterizing an entire order as having uniformly “small” or “large” population sizes may oversimplify the substantial variation within orders. It is also possible that our dataset, comprising only 12 orders, lacks the statistical power to detect such a relationship. Finally, there may truly be no connection between population size and chromosomal rearrangement rates. Of these possibilities, the oversimplification of population size within orders seems the most likely explanation, as we observed strong, consistent patterns when accounting for finer-scale variation.

We did find a significant relationship between population size and chromosomal fission and fusion rates when analyzing all bird species collectively. Specifically, we found that larger populations exhibited higher rates of fissions and fusions compared to smaller populations, suggesting that chromosomal fissions and fusions are significant evolutionary mechanisms that contribute to the adaptation of birds, and that selection is the dominant driver of chromosome number changes across birds. Our results align with findings by Hooper and Price (2015), who observed higher fixation rates of chromosomal inversions in genetically diverse, larger mainland Estrildid finches compared to smaller island populations, suggesting positive selection as the dominant mechanism driving fixation of intrachromosomal variants in birds. Our finding does conflict or add nuance to a previous hypothesis that natural selection acted primarily to maintain stasis in bird karyotypes (Zhang 2018). Instead, we would argue that while natural selection certainly does act as a sieve and limit the number and type of structural mutations that can fix in populations, it also serves as the primary force in driving disparification of chromosome number across birds.

### Fusions vs Fissions

We found that in our analysis of all birds rates of fusions were higher than rates of fissions. This pattern appears to be strongly driven by the signal from Passeriformes. In our order level analyses we found that Passeriformes was the only order of birds to exhibit unequal rates of chromosomal fissions and fusions, with significantly higher rates of fusions than fissions. Within passerines migratory lineages had higher fusion rates than sedentary lineages. This pattern may be explained by several factors. First, migratory species often maintain larger effective population sizes (Winker and Delmore 2023), which reduces the effects of drift. In such populations, chromosomal fusions that might otherwise be lost to drift in smaller, more isolated sedentary populations can persist, potentially accumulating over time. Second, selective pressures associated with long-distance navigation could favor chromosomal fusions if they enhance gene regulation or create co-adapted gene complexes that are advantageous in divergent environments (Stebbins 1971; Liu et al. 2022). Finally, fusions may confer energy efficiency by reducing the total number of chromosomes, which could lower the energetic costs of cellular processes, a potential advantage for species engaged in metabolically-intense activities like long-distance flight for migration (Waltari and Edwards 2002; Wright et al. 2014; Gregory 2018).

### Microchromosomes vs Macrochromosomes

Our findings show that the number of microchromosomes is negatively correlated with the number of macrochromosomes (**Figure 4**). This suggests that to some extent chromosome number can stay constant with only the partitioning between microchromosomes and macrochromosomes changing. This difference in partitioning could be due to biological reality (e.g., size expansion of microchromosome to such an extent that cytogeneticists classify them as microchromosomes) or due to differences in the heuristics that cytogeneticists use to classify chromosomes as microchromosomes or macrochromosomes as discussed in the introduction. In contrast if we look at the species that occur off the primary diagonal in **Figure 4**, we see species where fusions or fissions have changed the count of one or both size classes of chromosomes.

We observed a significant positive correlation between the evolutionary rates of microchromosomes and macrochromosomes across birds. This finding challenges the traditional view that microchromosomes evolve independently of macrochromosomes in birds (Damas et al. 2018; O’Connor et al. 2019). Instead, it suggests that shared mechanisms drive chromosomal rearrangements regardless of chromosome size. This could be mediated by the genomic architecture of evolutionary breakpoint regions (EBRs) that can span both microchromosomes and macrochromosomes (Claeys et al. 2023).

### Conclusion

Our study has several important limitations that should be acknowledged. First, the distinction between germline-restricted chromosomes (GRCs) and standard chromosomes was not explicitly addressed. GRCs are unique chromosomes found in germline cells but absent in somatic cells. They are known to occur in all Passeriformes but not other birds (Torgasheva et al. 2019). Not accounting for this variation could influence our rate calculations in Passeriformes. Secondly, the definition of microchromosomes is not universally standardized, and differences across studies biases among clades could lead to inaccuracies in our rate estimates.

The part of the Wright-Fisher debate in evolutionary biology centers around the relative importance of genetic drift and natural selection in shaping genetic variation and evolutionary trajectories (Provine 1992). This debate can be extended to the evolution of chromosome number across different clades, where some groups exhibit patterns consistent with Wrightian dynamics, dominated by genetic drift, while others show Fisherian characteristics, where selection plays a predominant role. In clades such as Coleoptera and Carnivora chromosome number evolution appears to be dominated by genetic drift (Blackmon et al. 2019, 2024; Jonika et al. 2024). Conversely, in birds chromosome number evolution appears to be dominated by natural selection.

## Acknowledgements

We thank the Sterling C. Evans Library at Texas A&M University for their assistance in obtaining primary sources. We also acknowledge the avian cytogenetics community for their dedication to fostering open and collaborative research.

## Data Availability Statement

The data as well all analysis code underlying this article are available in GitHub repository, at https://github.com/JamieAlfieri/bird.chroms.

---

**Supplemental Figure 1.**
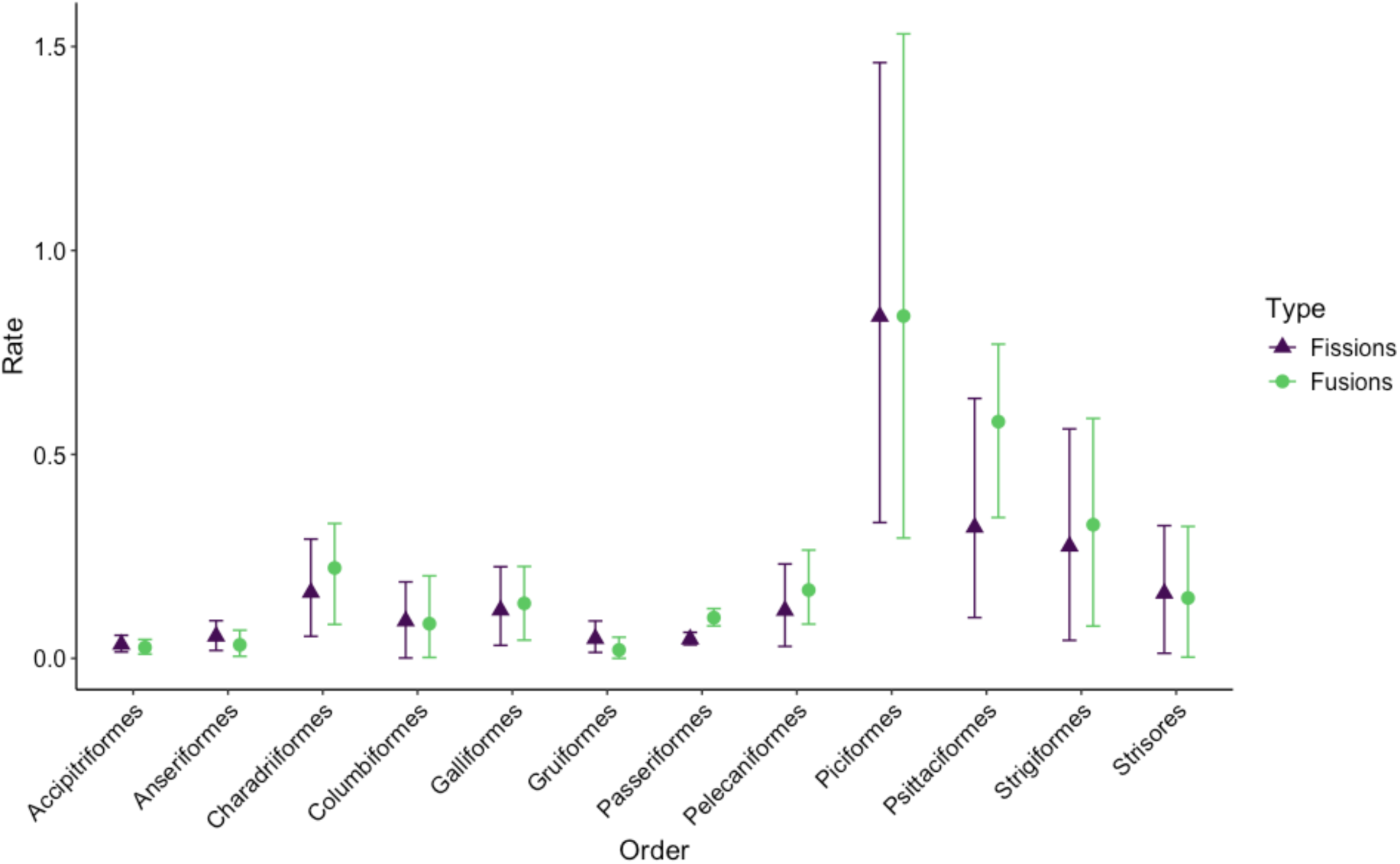
Chromosome fission and fusion rates for 12 bird orders. Bars represent the 95% credible interval.

**Supplemental Figure 2.**
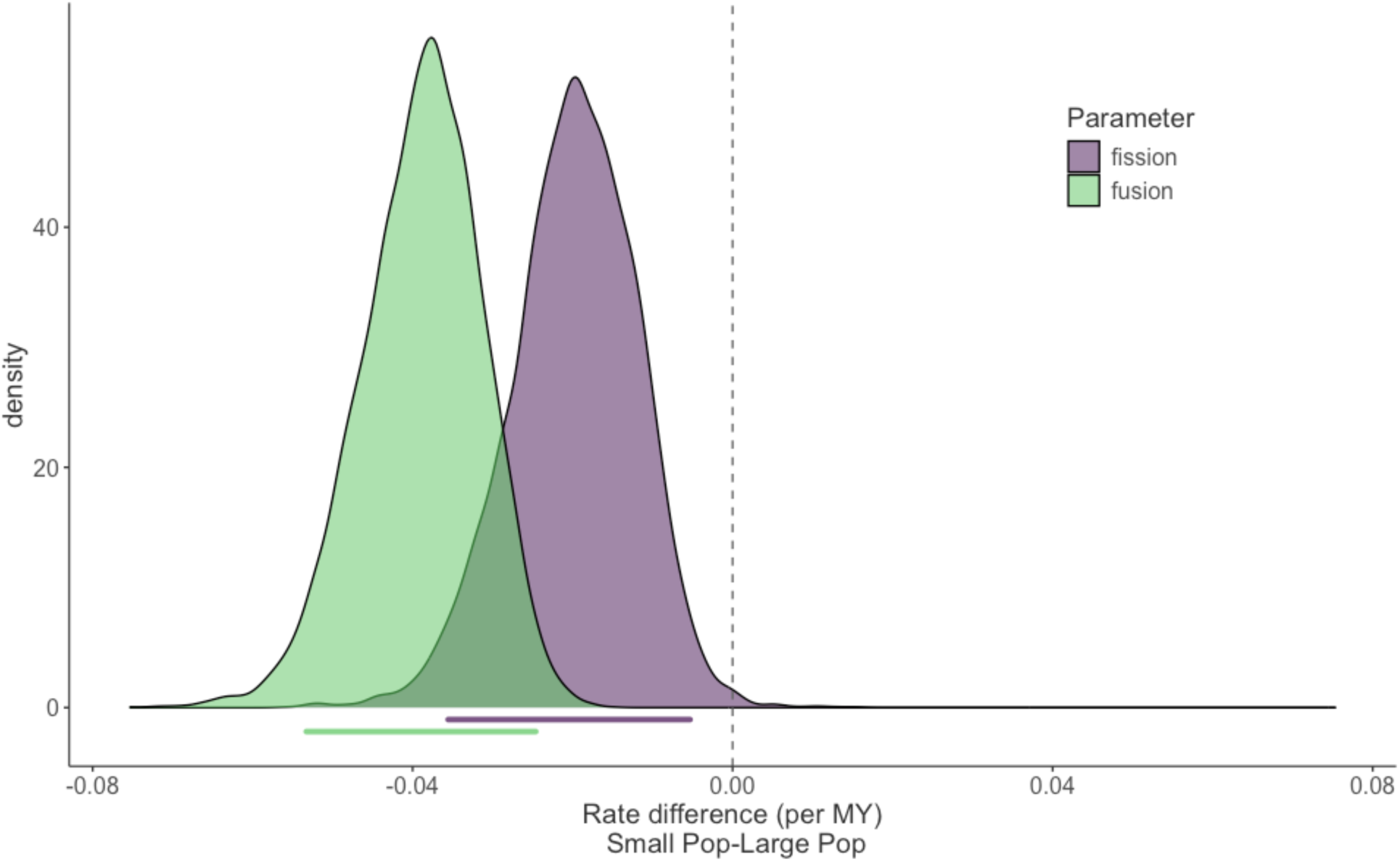
Chromosome evolution in small population size and large population size lineages excluding high tip rate taxa. Each curve shows the distribution of the rate difference in small population size and large population size lineages. The bars below the curve indicate the 95% credible interval for the rate differences.

**Supplemental Figure 3.**
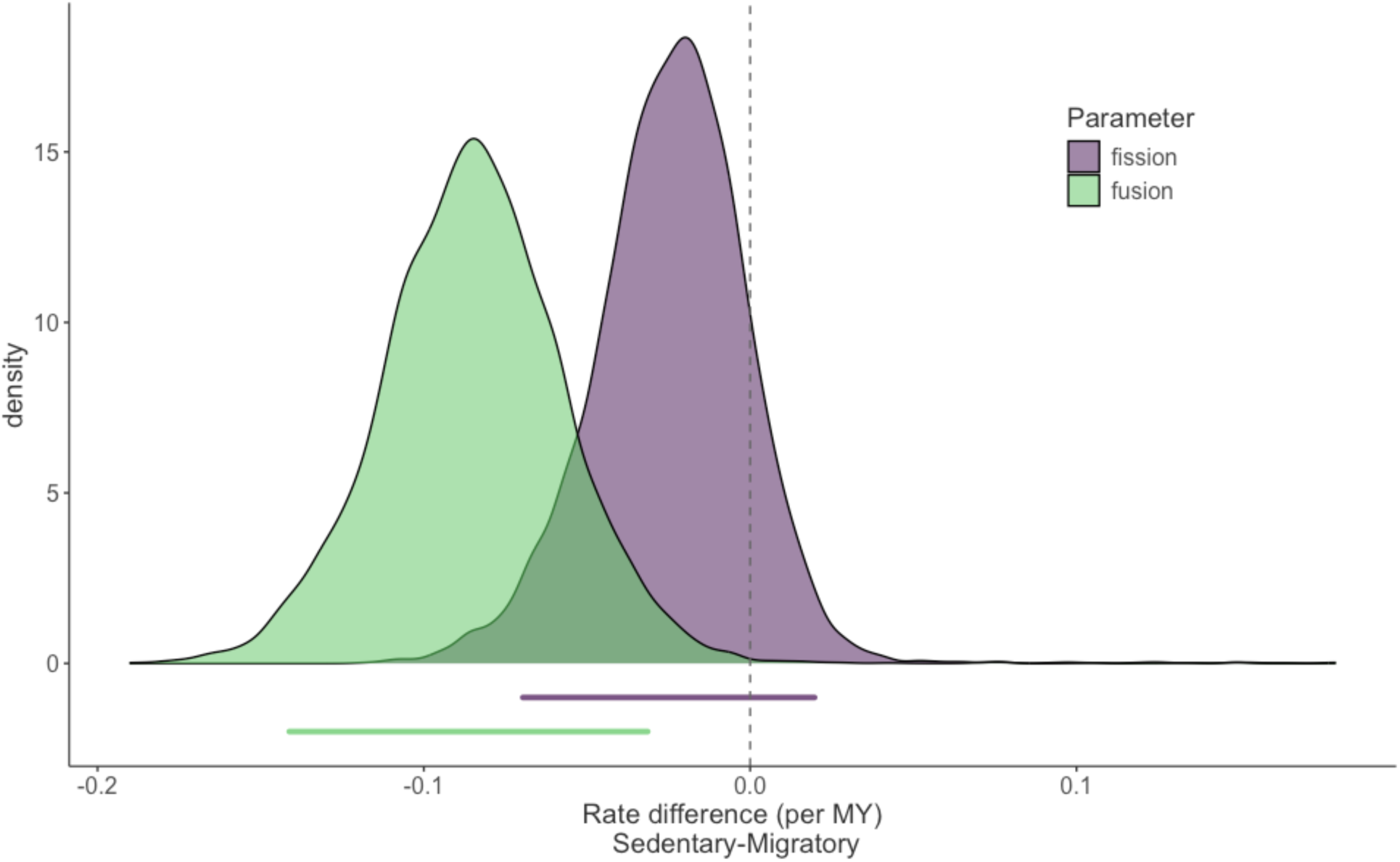
Chromosome evolution in migratory and sedentary Passeriformes. Each curve shows the distribution of the rate difference in sedentary and migratory lineages within Passeriformes. The bars below the curve indicate the 95% credible interval for the rate differences.

## References

Alfieri, J. M., M. M. Jonika, J. N. Dulin, and H. Blackmon. 2023. Tempo and mode of genome structure evolution in insects. Genes 14:336. MDPI AG.

Blackmon, H., M. Chin, and M. Jonika. 2023. chromePlus: Analysis of Chromosome Number Evolution and Binary Traits.

Blackmon, H., and J. P. Demuth. 2015. The fragile Y hypothesis: Y chromosome aneuploidy as a selective pressure in sex chromosome and meiotic mechanism evolution. Bioessays 37:942–950.

Blackmon, H., N. B. Hardy, and L. Ross. 2015. The evolutionary dynamics of haplodiploidy: Genome architecture and haploid viability. Evolution 69:2971–2978.

Blackmon, H., M. M. Jonika, J. M. Alfieri, L. Fardoun, and J. P. Demuth. 2024. Drift drives the evolution of chromosome number I: The impact of trait transitions on genome evolution in Coleoptera. J. Hered. 115:173–182. Oxford University Press (OUP).

Blackmon, H., J. Justison, I. Mayrose, and E. E. Goldberg. 2019. Meiotic drive shapes rates of karyotype evolution in mammals. Evolution 73:511–523. Wiley.

Britton-Davidian, J., H. Sonjaya, J. Catalan, and G. Cattaneo-Berrebi. 1990. Robertsonian heterozygosity in wild mice: fertility and transmission rates in Rb(16.17) translocation heterozygotes. Genetica 80:171–174. Springer Nature.

Brüniche-Olsen, A., K. F. Kellner, J. L. Belant, and J. A. DeWoody. 2021. Life-history traits and habitat availability shape genomic diversity in birds: implications for conservation. Proc. Biol. Sci. 288:20211441.

Burt, D. W. 2002. Origin and evolution of avian microchromosomes. Cytogenet. Genome Res. 96:97–112. S. Karger AG.

Bush, G. L., S. M. Case, A. C. Wilson, and J. L. Patton. 1977. Rapid speciation and chromosomal evolution in mammals. Proc. Natl. Acad. Sci. U. S. A. 74:3942–3946.

Callaghan, C. T., S. Nakagawa, and W. K. Cornwell. 2021. Global abundance estimates for 9,700 bird species. Proc. Natl. Acad. Sci. U. S. A. 118:e2023170118. Proceedings of the National Academy of Sciences.

Claeys, J., M. N. Romanov, and D. K. Griffin. 2023. Integrative comparative analysis of avian chromosome evolution by in-silico mapping of the gene ontology of homologous synteny blocks and evolutionary breakpoint regions. Genetica 151:167–178. Springer Science and Business Media LLC.

Damas, J., J. Kim, M. Farré, D. K. Griffin, and D. M. Larkin. 2018. Reconstruction of avian ancestral karyotypes reveals differences in the evolutionary history of macro- and microchromosomes. Genome Biol. 19:155. Springer Science and Business Media LLC.

Degrandi, T. M., S. A. Barcellos, A. L. Costa, A. D. V. Garnero, I. Hass, and R. J. Gunski. 2020. Introducing the Bird Chromosome Database: An overview of cytogenetic studies in birds. Cytogenet. Genome Res. 160:199–205. S. Karger AG.

de Oliveira Furo, I. R. Kretschmer, M. S. Dos Santos, C. A. de Lima Carvalho, R. J. Gunski, P. C. M. O’Brien, M. A. Ferguson-Smith, M. B. Cioffi, and E. H. C. de Oliveira. 2017. Chromosomal Mapping of Repetitive DNAs in Myiopsitta monachus and Amazona aestiva (Psittaciformes, Psittacidae) with Emphasis on the Sex Chromosomes. Cytogenet. Genome Res. 151:151–160. S. Karger AG.

Ebied, A. M., H. A. Hassan, A. H. Abu Almaaty, and A. E. Yaseen. 2005. Karyotypic characterization of ten species of birds. Cytologia (Tokyo) 70:181–194. International Society of Cytology.

Ellegren, H. 2010. Evolutionary stasis: the stable chromosomes of birds. Trends Ecol. Evol. 25:283–291. Elsevier BV.

Eo, S. H., J. M. Doyle, and J. A. DeWoody. 2011. Genetic diversity in birds is associated with body mass and habitat type: Microsatellite diversity in birds. J. Zool. (1987) 283:220–226. Wiley.

FitzJohn, R. G. 2012. Diversitree: comparative phylogenetic analyses of diversification in R. Methods in Ecology and Evolution 3:1084–1092. John Wiley & Sons, Ltd.

Freyman, W. A., and S. Höhna. 2018. Cladogenetic and anagenetic models of chromosome number evolution: A Bayesian model averaging approach. Syst. Biol. 67:195–215.

Gaston, K. J., and T. M. Blackburn. 1996. Global scale macroecology: Interactions between population size, geographic range size and body size in the Anseriformes. J. Anim. Ecol. 65:701. JSTOR.

Glick, L., and I. Mayrose. 2014. ChromEvol: assessing the pattern of chromosome number evolution and the inference of polyploidy along a phylogeny. Molecular biology and evolution 31:1914–1922. academic.oup.com.

Goldschmidt, B., D. M. Nogueira, D. W. Monsores, and L. M. Souza. 1997. Chromosome study in two Aratinga species (A. guarouba and A. acuticaudata) (Psittaciformes). Rev. Bras. Genet. 20:659–662. FapUNIFESP (SciELO).

Gregory, T. R. 2018. Genome size and brain cell density in birds. Can. J. Zool. 96:379–382. Canadian Science Publishing.

Griffin, D. K., L. B. W. Robertson, H. G. Tempest, and B. M. Skinner. 2007. The evolution of the avian genome as revealed by comparative molecular cytogenetics. Cytogenet. Genome Res. 117:64–77. Cytogenet Genome Res.

Hassan, H. A. 1998. Karyological studies on six species of birds. Cytologia (Tokyo) 63:349–363. International Society of Cytology.

Hooper, D. M., and T. D. Price. 2015. Rates of karyotypic evolution in Estrildid finches differ between island and continental clades. Evolution 69:890–903. Wiley.

International Chicken Genome Sequencing Consortium. 2004. Sequence and comparative analysis of the chicken genome provide unique perspectives on vertebrate evolution. Nature 432:695–716. Springer Science and Business Media LLC.

Jonika, M. M., M. Chin, N. Anderson, R. H. Adams, J. P. Demuth, and B. H. 2023. EvoBiR: Tools for comparative analyses and teaching evolutionary biology.

Jonika, M. M., K. T. Wilhoit, M. Chin, A. Arekere, and H. Blackmon. 2024. Drift drives the evolution of chromosome number II: The impact of range size on genome evolution in Carnivora. J. Hered. 115:524–531. Oxford University Press (OUP).

Kaul, D., and H. A. Ansari. 1978. Chromosome studies in three species of Piciformes (Aves). Genetica 48:193–196. Springer Nature.

King, M. 1995. Species evolution: The role of chromosome change. Cambridge University Press, Cambridge, England.

Kretschmer, R., M. S. de Souza, I. de O. Furo, M. N. Romanov, R. J. Gunski, A. D. V. Garnero, T. R. O. de Freitas, E. H. C. de Oliveira, R. E. O’Connor, and D. K. Griffin. 2021. Interspecies chromosome mapping in Caprimulgiformes, Piciformes, Suliformes, and Trogoniformes (Aves): Cytogenomic insight into microchromosome organization and karyotype evolution in birds. Cells 10:826. MDPI AG.

Lande, R. 1979. Effective Deme sizes during long-term evolution estimated from rates of chromosomal rearrangement. Evolution 33:234–251. Wiley.

Li, J., J. Zhang, J. Liu, Y. Zhou, C. Cai, L. Xu, X. Dai, S. Feng, C. Guo, J. Rao, K. Wei, E. D. Jarvis, Y. Jiang, Z. Zhou, G. Zhang, and Q. Zhou. 2021. A new duck genome reveals conserved and convergently evolved chromosome architectures of birds and mammals. Gigascience 10:giaa142. Oxford University Press (OUP).

Liu, Z., M. Roesti, D. Marques, M. Hiltbrunner, V. Saladin, and C. L. Peichel. 2022. Chromosomal fusions facilitate adaptation to divergent environments in threespine stickleback. Mol. Biol. Evol. 39:msab358. Oxford University Press (OUP).

Mackintosh, A., R. Vila, S. H. Martin, D. Setter, and K. Lohse. 2023. Do chromosome rearrangements fix by genetic drift or natural selection? Insights from Brenthis butterflies. Mol. Ecol., doi: 10.1111/mec.17146. Wiley.

Mayrose, I., M. S. Barker, and S. P. Otto. 2010. Probabilistic models of chromosome number evolution and the inference of polyploidy. Syst. Biol. 59:132–144.

McTavish, E. J., J. A. Gerbracht, M. T. Holder, M. J. Iliff, D. Lepage, P. Rasmussen, B. Redelings, L. L. Sanchez Reyes, and E. T. Miller. 2024. A complete and dynamic tree of birds.

Muller, H. J. 1930. Types of visible variations induced by X-rays in Drosophila. Journal of genetics 22:299–334.

O’Connor, R. E., L. Kiazim, B. Skinner, G. Fonseka, S. Joseph, R. Jennings, D. M. Larkin, and D. K. Griffin. 2019. Patterns of microchromosome organization remain highly conserved throughout avian evolution. Chromosoma 128:21–29. Springer Science and Business Media LLC.

O’Connor, R. E., R. Kretschmer, M. N. Romanov, and D. K. Griffin. 2024. A bird’s-eye view of chromosomic evolution in the Class Aves. Cells 13:310. MDPI AG.

O’Connor, R. E., M. N. Romanov, L. G. Kiazim, P. M. Barrett, M. Farré, J. Damas, M. Ferguson-Smith, N. Valenzuela, D. M. Larkin, and D. K. Griffin. 2018. Reconstruction of the diapsid ancestral genome permits chromosome evolution tracing in avian and non-avian dinosaurs. Nat. Commun. 9:1883. Nature Publishing Group.

Petitpierre, E. 1987. Why beetles have strikingly different rates of chromosomal evolution. Elytron 1:25–32.

Pichugin, A. M., S. A. Galkina, A. A. Potekhin, E. O. Punina, M. S. Rautian, and A. V. Rodionov. 2001. Estimation of the Minimal Size of Chicken Gallus gallus domesticusMicrochromosomes via Pulsed-Field Electrophoresis. Russ. J. Genet. 37:535–538. Springer Nature.

Provine, W. B. 1992. The R. a. fisher—Sewall wright controversy. Pp. 201–229 in Boston Studies in the Philosophy of Science. Springer Netherlands, Dordrecht.

Qumsiyeh, M., and E. Handal. 2022. Adaptive nature of chromosome variation in placental mammals and applicability to domestication and invasiveness. Hystrix It. J. Mamm. 33:103–107. Associazione Teriologica Italiana.

Rabosky, D. L., and E. E. Goldberg. 2015. Model inadequacy and mistaken inferences of trait-dependent speciation. Syst. Biol. 64:340–355.

Ratomponirina, C. 1988. Synaptonemal complexes in Robertsonian translocation heterozygous in lemurs. Kew Chromosomes. HMSO Publ. Centre.

R Core Team. 2024. R: A Language and Environment for Statistical Computing.

Revell, L. J. 2012. phytools: An R package for phylogenetic comparative biology (and other things).

Rodionov, A. V., L. A. Chel’sheva, I. V. Soloveĭ, and I. A. Miakoshina. 1992. Chiasma distribution in the lampbrush chromosomes of the chicken Gallus gallus domesticus: hot spots of recombination and their possible role in proper dysjunction of homologous chromosomes at the first meiotic division. Genetika 28:151–160. Genetika.

Romanov, M. N., M. Farré, P. E. Lithgow, K. E. Fowler, B. M. Skinner, R. O’Connor, G. Fonseka, N. Backström, Y. Matsuda, C. Nishida, P. Houde, E. D. Jarvis, H. Ellegren, D. W. Burt, D. M. Larkin, and D. K. Griffin. 2014. Reconstruction of gross avian genome structure, organization and evolution suggests that the chicken lineage most closely resembles the dinosaur avian ancestor. BMC Genomics 15:1060. Springer Science and Business Media LLC.

Skinner, B. M., and D. K. Griffin. 2012. Intrachromosomal rearrangements in avian genome evolution: evidence for regions prone to breakpoints. Heredity (Edinb.) 108:37–41. Springer Science and Business Media LLC.

Smith, J., C. K. Bruley, I. R. Paton, I. Dunn, C. T. Jones, D. Windsor, D. R. Morrice, A. S. Law, J. Masabanda, A. Sazanov, D. Waddington, R. Fries, and D. W. Burt. 2000. Differences in gene density on chicken macrochromosomes and microchromosomes. Anim. Genet. 31:96–103. Wiley.

Sokoloff, A. 1977. The biology of Tribolium with special emphasis on genetic aspects. Volume 3. cabidigitallibrary.org.

Stebbins, G. L. 1971. Chromosomal evolution in higher plants. Edward Arnold, London. Stebbins, G. L. 1958. *Longevity, habitat, and release of genetic variability in the higher plants*. Cold Spring Harb. Symp. Quant. Biol. 23:365–378.

Tobias, J. A., C. Sheard, A. L. Pigot, A. J. M. Devenish, J. Yang, F. Sayol, M. H. C. Neate-Clegg, N. Alioravainen, T. L. Weeks, R. A. Barber, P. A. Walkden, H. E. A. MacGregor, S. E. I. Jones, C. Vincent, A. G. Phillips, N. M. Marples, F. A. Montaño-Centellas, V. Leandro-Silva, S. Claramunt, B. Darski, B. G. Freeman, T. P. Bregman, C. R. Cooney, E. C. Hughes, E. J. R. Capp, Z. K. Varley, N. R. Friedman, H. Korntheuer, A. Corrales-Vargas, C. H. Trisos, B. C. Weeks, D. M. Hanz, T. Töpfer, G. A. Bravo, V. Remeš, L. Nowak, L. S. Carneiro, A. J. Moncada R, B. Matysioková, D. T. Baldassarre, A. Martínez-Salinas, J. D. Wolfe, P. M. Chapman, B. G. Daly, M. C. Sorensen, A. Neu, M. A. Ford, R. J. Mayhew, L. Fabio Silveira, D. J. Kelly, N. N. D. Annorbah, H. S. Pollock, A. M. Grabowska-Zhang, J. P. McEntee, J. Carlos T Gonzalez, C. G. Meneses, M. C. Muñoz, L. L. Powell, G. A. Jamie, T. J. Matthews, O. Johnson, G. R. R. Brito, K. Zyskowski, R. Crates, M. G. Harvey, M. Jurado Zevallos, P. A. Hosner, T. Bradfer-Lawrence, J. M. Maley, F. G. Stiles, H. S. Lima, K. L. Provost, M. Chibesa, M. Mashao, J. T. Howard, E. Mlamba, M. A. H. Chua, B. Li, M. I. Gómez, N. C. García, M. Päckert, J. Fuchs, J. R. Ali, E. P. Derryberry, M. L. Carlson, R. C. Urriza, K. E. Brzeski, D. M. Prawiradilaga, M. J. Rayner, E. T. Miller, R. C. K. Bowie, R.-M. Lafontaine, R. P. Scofield, Y. Lou, L. Somarathna, D. Lepage, M. Illif, E. L. Neuschulz, M. Templin, D. M. Dehling, J. C. Cooper, O. S. G. Pauwels, K. Analuddin, J. Fjeldså, N. Seddon, P. R. Sweet, F. A. J. DeClerck, L. N. Naka, J. D. Brawn, A. Aleixo, K. Böhning-Gaese, C. Rahbek, S. A. Fritz, G. H. Thomas, and M. Schleuning. 2022. AVONET: morphological, ecological and geographical data for all birds. Ecol. Lett. 25:581–597.

Torgasheva, A. A., L. P. Malinovskaya, K. S. Zadesenets, T. V. Karamysheva, E. A. Kizilova, E. A. Akberdina, I. E. Pristyazhnyuk, E. P. Shnaider, V. A. Volodkina, A. F. Saifitdinova, S. A. Galkina, D. M. Larkin, N. B. Rubtsov, and P. M. Borodin. 2019. Germline-restricted chromosome (GRC) is widespread among songbirds. Proc. Natl. Acad. Sci. U. S. A. 116:11845–11850. Proceedings of the National Academy of Sciences.

Waltari, E., and S. V. Edwards. 2002. Evolutionary dynamics of intron size, genome size, and physiological correlates in archosaurs. Am. Nat. 160:539–552. University of Chicago Press.

Waters, P. D., H. R. Patel, A. Ruiz-Herrera, L. Álvarez-González, N. C. Lister, O. Simakov, T. Ezaz, P. Kaur, C. Frere, F. Grützner, A. Georges, and J. A. M. Graves. 2021. Microchromosomes are building blocks of bird, reptile, and mammal chromosomes. Proc. Natl. Acad. Sci. U.S.A. 118:e2112494118. National Academy of Sciences.

White, M. J. D. 1973. Animal Cytology and Evolution. Cambridge University Press.

Wilson, A. C., G. L. Bush, S. M. Case, and M. C. King. 1975. Social structuring of mammalian populations and rate of chromosomal evolution. Proc. Natl. Acad. Sci. U. S. A. 72:5061–5065.

Winker, K., and K. Delmore. 2023. Seasonally migratory songbirds have different historic population size characteristics than resident relatives.

Wright, N. A., T. R. Gregory, and C. C. Witt. 2014. Metabolic “engines” of flight drive genome size reduction in birds. Proc. Biol. Sci. 281:20132780. The Royal Society.

Zenil-Ferguson, R., J. M. Ponciano, and J. G. Burleigh. 2017. Testing the association of phenotypes with polyploidy: An example using herbaceous and woody eudicots. Evolution 71:1138–1148.

Zhang, G. 2018. The bird’s-eye view on chromosome evolution. Genome Biol. 19:201. Springer Science and Business Media LLC.

